# Large Language Models Predict Human Social Behavior via Interpretable Mechanisms

**DOI:** 10.64898/2026.09.06.749667

**Authors:** Yingying Ye, Youle Fang, Xiaoxue Gao, Vered Kurtz-David, Andreas Hula, Chun Fan, Tingting Long, Bo Shen

**Author notes:** Corresponding Author: Tingting Long, Bo Shen. **Co-first authors:** Yingying Ye and Youle Fang contributed equally to this work.

## Abstract

The development of large language models (LLMs) offers promising opportunities for predicting human behavior across diverse contexts. However, most prior work has emphasized behavioral imitation, with limited attention to transparent or interpretable models of the cognitive mechanisms underlying human decisions. In this study, we introduce MindEvolve, an autonomous workflow designed to predict behavior in social interactions by generating interpretable symbolic models of cognition. We systematically evaluate the modeling capabilities of multiple LLMs across a battery of socioeconomic games covering four core domains of social cognition: economic preferences, social preferences, social reasoning (theory of mind), and recursive planning. The symbolic models proposed by the LLMs are assessed by expert human evaluators for interpretability and theoretical coherence. Our results show that while most LLMs robustly capture economic and social preferences in relatively simple strategic settings—and in some cases generate novel models that integrate broader knowledge than those proposed by human experts—their capacity to model more complex psychological processes remains limited. However, a subset of state-of-the-art models demonstrates promising performance in capturing higher-order reasoning processes such as theory of mind and recursive planning. Together, these findings highlight the emerging potential of LLMs not only to imitate behavior, but to generate interpretable, mechanistic accounts of human social decision-making. Our results provide a roadmap for advancing LLM-based cognitive modeling toward human-expert-level theory construction.

**Significance Statement:** Large language models can increasingly predict human behavior, but whether they can construct interpretable models of the cognitive mechanisms underlying behavior remains unclear. We introduce MindEvolve, a framework that uses large language models to autonomously build symbolic models of human social decision-making. Across tasks involving economic and social preferences, theory of mind, and repeated trust interactions, these models generated transparent, expert-evaluable cognitive models and sometimes proposed theoretically meaningful strategies beyond existing approaches. Although complex social reasoning and long-horizon planning remain challenging, advanced models already show emerging abilities to capture higher-order cognitive processes. These findings suggest that large language models may become powerful scientific tools for constructing and discovering theories of human cognition.

## Introduction

The development of large language models (LLMs) presents unprecedented opportunities to understand and predict human behavior in non-traditional ways. An increasing number of studies precisely predict human behavior across a variety of psychological paradigms using LLMs (Cheung et al., 2025; Kosinski, 2024; Mei et al., 2024; Strachan et al., 2024). By fine-tuning LLMs on large-scale behavioral datasets, research even shows that a single artificial agent can predict human participants’ behavior across a wide range of social and psychological tasks (Binz et al., 2025). The development of AI-driven approaches to predicting human behavior in social contexts presents exciting opportunities and has the potential to advance social AI agents capable of predicting human social interactions.

However, it has been criticized that the internal mechanisms underlying LLMs’ ability to generate human-like behaviors may still rely primarily on statistical patterns (Gurney et al., 2024; Trott et al., 2023). Most LLMs generated behavior is largely based on a mechanism of imitation while lacking an interpretable understanding of how the LLMs capture human behavior. How exactly do LLMs understand human behavior? It remains challenging to reveal whether LLMs capture the underlying structure of social tasks and the psychological processes that drives human behavior.

Modeling human social interactions is highly complex (Gao et al., 2024; Wang et al., 2024), since social scenarios require LLMs to recognize and interpret subtle emotional cues (Sabour et al., 2024), infer latent intentions, understand nuanced social contexts (Zhang et al., 2025), and assess the truthfulness of information (Xu et al., 2025). Using approaches from cognitive science and psychology, many aspects of human behavior can be quantified and described using symbolic models. Computational and symbolic modeling, through the construction of precise mathematical models or equations (Wilson & Collins, 2019), offers a promising methodological approach for examining how LLMs understand and predict human cognition and behavior.

LLMs have been applied to explore symbolic models in computer science and mathematics, yielding novel algorithms and solutions, notable examples including AlphaGeometry (Trinh et al., 2024) and FunSearch (Romera-Paredes et al., 2024). The FunSearch is a representative framework for symbolic modeling that has made new discoveries on open mathematical problems (Romera-Paredes et al., 2024). Its core principle is to have LLMs generate candidate models from problem prompts, which are then progressively optimized through an automated evolutionary process. The framework has been extended to modeling human cognition, for example, in CogFunSearch (Castro et al., 2025). LLMs autonomously extract structure from vast corpora and datasets, generating rules and symbolic representations that may fundamentally reshape the paradigm of computational modeling.

Employing LLMs for computational modeling of social interactions offers significant advantages and addresses key gaps in previous research. Previous studies have primarily evaluated the social abilities of LLMs by measuring output features related to human behavior (Mei et al., 2024) or by simulating social interaction scenarios (Akata et al., 2025; Binz et al., 2025; Liu et al., 2024; Xu et al., 2025; Zhang et al., 2025). By contrast, symbolic models enable accurate quantitative predictions of human social behavior through transparent processes with strong interpretability, allowing tracing each step in the LLMs’ reasoning. Leveraging the broad, cross-disciplinary knowledge embedded in LLMs, the current study tests their ability to construct symbolic models that predict human behavior across a variety of tasks and social contexts.

Due to the richness of social interaction contexts, we tested a range of tasks to cover a set of important psychological factors. Symbolic modeling for economic preferences in the literature shows that humans’ behavior often deviates from rational optimization, exhibiting behaviors like risk aversion and loss aversion when making economic decisions (Kahneman & Tversky, 1979). Traditionally, human experts build such symbolic models with specific parameters to capture risk aversion and loss aversion idiosyncrasies (Kahneman & Tversky, 1979). In addition to economic preferences, individuals often show other-regarding concerns beyond maximizing their own benefits (Fehr & Schmidt, 1999). Literature shows that these behaviors can be precisely captured and interpreted using symbolic models with weighting parameters between the welfares of one’s own and others (Charness & Dufwenberg, 2006; Fehr & Schmidt, 1999, 2006). To best test these psychological factors, we choose a one-shot socio-economic game named the Dictator Game (Gao et al., 2018; Sáez et al., 2015), with adaptation from its original version (Forsythe et al., 1994), where the decision maker trades off the economic payoffs between themselves and an anonymous partner under different social contexts. The game primarily reflects altruistic behavior, and its modeling requires consideration of both economic preferences (e.g., understanding the reward structure of the task to maximize one’s own payoff) and social preferences (e.g., other’s payoff and inequality aversion), and the social contexts.

Beyond that, theory of mind, the ability to consider others’ actions in one’s own decisions, plays a crucial role in comprehending and responding to complex social interactions (Lore et al., 2024; Gurney et al., 2024; Strachan et al., 2024). This skill is considered one of the key cognitive features of human social interactions (Devaine et al., 2017; Premack & Woodruff, 1978). Theory of mind exhibits a recursive nature (Devaine et al., 2017; Hula et al., 2021), as mentalizing about others’ thoughts and perspectives represents an advanced cognitive function possessed by only some mammals (Bennett et al., 2021). Symbolic models of theory of mind allow for precise quantification of participants’ key characteristics via parameterized representations (Hula et al., 2018; Reiter et al., 2023), making social reasoning processes transparent and interpretable while offering mechanistic accounts of the underlying system (Gao et al., 2024).

Recent studies have shown inconsistency about whether the state-of-the-art LLMs can perform well in a range of theory of mind tasks (Bubeck et al., 2023; Gandhi et al., 2023; Kosinski, 2024; Holterman & van Deemter, 2023; Moghaddam & Honey, 2023; Strachan et al., 2024; Trott et al., 2023; Shapira et al., 2024; Ullman, 2023). LLMs particularly fail in higher-order aspects where their performance does not surpass that of human experts (Chan et al., 2024; van Duijn et al., 2023; Jones et al., 2024; Kim et al., 2023; Sap et al., 2022). These inconsistencies may reflect certain limitations in LLMs’ understanding of the theory of mind. In dynamic interactions between players, however, LLMs usually fail to capture social interactions and social inference (Bubeck et al., 2023; Gao et al., 2024; Xu et al., 2025). To assess the ability of LMMs for modeling theory of mind, we use a game-theoretic paradigm named the “hide-and-seek” location game (Li & Camerer, 2022). This game requires consideration not only of the player’s payoff structure within the task context, but also of strategic interaction between two players, necessitating social reasoning about another agent. It therefore constitutes an ideal paradigm for studying theory of mind.

Beyond that, dynamic multi-trial social interactions may require additional recursive planning across many trials to maximize returns, other than theory-of-mind simulation. In an investment game named “Trust Game” (Berg et al., 1995; McCabe et al., 2001), the investor optimizes their investment amount to the trustees based on the return history of the trustees and potential returns in future trials to maximize their return (Charness & Dufwenberg, 2006; Hula et al., 2018; McCabe et al., 1998; Reiter et al., 2023). Thus, modeling recursive planning often requires iterative optimization processes, such as dynamic programming (Bellman, 1958), which introduces larger computational complexity than in one-shot games. We tested a multi-trial version of the Trust Game that involves both theory of mind and recursive planning over many trials between the investors and the trustees (Hula et al., 2018; Reiter et al., 2023).

At the time of our execution, we tested a variety of LLMs, including non-reasoning models, such as Deepseek V3, and reasoning models, such as Deepseek R1, to assess whether these LLMs are capable of modeling social behaviors across the four domains of psychological factors mentioned above, i.e., economic preferences, social preferences, theory of mind, and multi-trial planning. The goal is not to exhaustively compare across LLM versions. Instead, we aim to provide a broader view over versions of LLMs that have been trained in different manners.

For assessing the capacity of LLMs, we propose a workflow named MindEvolve (Fig. 1). Current workflows, like the ones mentioned (e.g., AlphaGeometry, FunSearch, and CogFunSearch), rely on unsupervised evolutionary search to retain high-scoring solutions and eliminate low-scoring ones, a process that typically consumes substantial computational resources and requires sampling tens or even hundreds of thousands of programs before identifying high-quality solutions (Castro et al., 2025). MindEvolve aims to enhance iteration efficiency at both the algorithmic and theoretical levels by introducing several strategies, including feedback loops, a self-reviewing process, and multi-objective evolutionary search algorithms. Building on the research of social and cognitive science, we systematically assess LLMs’ performance in their goodness of fit to human’s behavioral data, as well as the identifiability of the parameters and their interpretability. Human evaluators with expertise in these specific games were invited to assess the symbolic models generated by the LLMs.

**Fig. 1.**
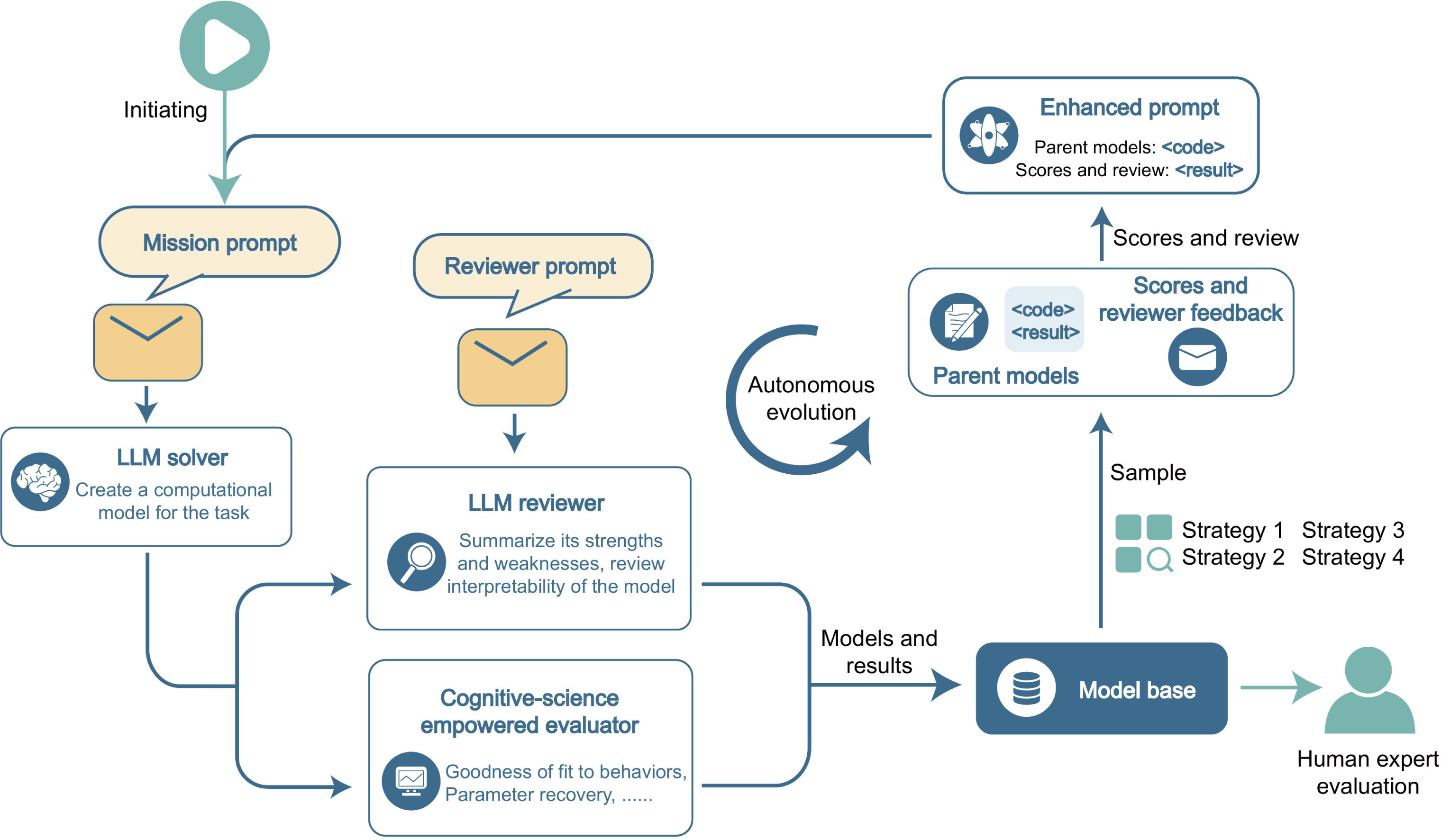
The workflow consisted of a cognitive-science empowered autonomous evolution loop and human expert evaluation. The autonomous evaluation loop iteratively improves the proposed computational models to explain human players’ behaviors. The loop is initiated by a mission prompt that specified the game, defined the players’ response variables, and set the goal as building a computational model to explain the players’ behaviors. The LLM solver proposes 20 alternative models from each generation. Cognitive science approaches are used to evaluate the proposed models’ performance in explaining the data, such as goodness of fit and parameter identifiability. Other than that, we invited another LLM agent to review the proposed models for their psychological plausibility and the interpretability of each parameter. The models and their evaluation results were stored in the model base and subsequently selected as parent models to improve the next generation. During evolution, we selected parent models using four strategies, ranging from random selection to selecting the top-scoring models (Methods). An enhanced prompt was used to improve the models for the next generations. After five generations, the models’ performance saturated. The best model of each game from the autonomous loop was evaluated by two human experts.

Our results indicate that while most LLMs can effectively model basic social and economic preferences, only a few of the most recent models (e.g., Claude 3.7 Sonnet and Gemini 2.5 Pro) exhibit emerging capabilities in modeling more complex constructs such as theory of mind and planning. These findings inform the development of AI systems capable of capturing the social and psychological determinants of human behavior in an interpretable way.

## Results

### The workflow of MindEvolve

The MindEvolve workflow included an autonomous loop within the LLM agents to self-evolve and a transparent output that enables human evaluation (Fig. 1). The goal of the LLM agents was to predict human players’ behaviors in the game. To improve the transparency of our approach, we asked the LLM agents (named LLM solver) to explicitly write a Python-based computational model to predict the behaviors, rather than generating behaviors implicitly from their hidden-layer embeddings. This approach made every step of the model’s prediction transparent and tractable.

To facilitate this process, we built an autonomous loop using a evolutionary algorithm to evolve the model over generations (Methods). The autonomous evolution loop was initiated from the LLM solver, which received a detailed description of the task, definitions of the input and response variables, and the required model interface, but participant-level choices or trial-by-trial behavioral records were included in its prompt. Cognitive science approaches were used to evaluate the model’s goodness-of-fit to the behavioral data by penalizing the number of parameters using the Bayesian information criterion (BIC). Crucially, to improve the identifiability of each parameter to reduce parameter dependencies when capturing the psychological factors. Thus, we assessed parameter recovery by quantifying the dependency between each parameter pair using *R*^2^.

Meanwhile, to develop computational models that best predict behavior while maintaining high interpretability for human evaluation, we invited a second LLM agent (LLM reviewer) into the autonomous loop to score the psychological interpretability of the proposed models and to provide qualitative feedback in the form of review scores for each parameter and a summary of the model’s strengths and weaknesses. The review score was combined with other model evaluation metrics and jointly fed to the LLM solver for refinement over iterations. Accordingly, the autonomous loop selected the highest-scoring candidate under the predefined evaluation criteria, which included in-sample fit, parameter recovery, and interpretability.

To improve the efficiency of model evolution, a commonly used ablation study (Meyes, 2019) from machine learning was applied to identify the key components of the proposed models and remove unnecessary components. To do that, we proposed parallelizing model evolution and storing the 20 models produced at each iteration in the model base (Fig. 1). Two models from every iteration were selected as parents for subsequent iteration refinement. For this process, we tested four evolutionary strategies for parent model selection and improvement feedback (Methods). Strategies #1 and #2 randomly sampled parent models for the next generation. Strategies #3 and #4 sampled from the model base sorted by performance and employ a structured pairing strategy, where parent models were selected according to their rankings in BIC scores and LLM reviewer scores (see details in Methods). Among the four tested ablation strategies, we consistently revealed the strategy (strategy #4) that paired two parent models with top scores in both model fitting and reviewing interpretability can effectively improve both goodness of fit and model interpretability. To illustrate this process, the ablation study resulted from DeepSeek R1 (250528) were visualized for all tasks in Figs. 2-4.

**Figure 2.**
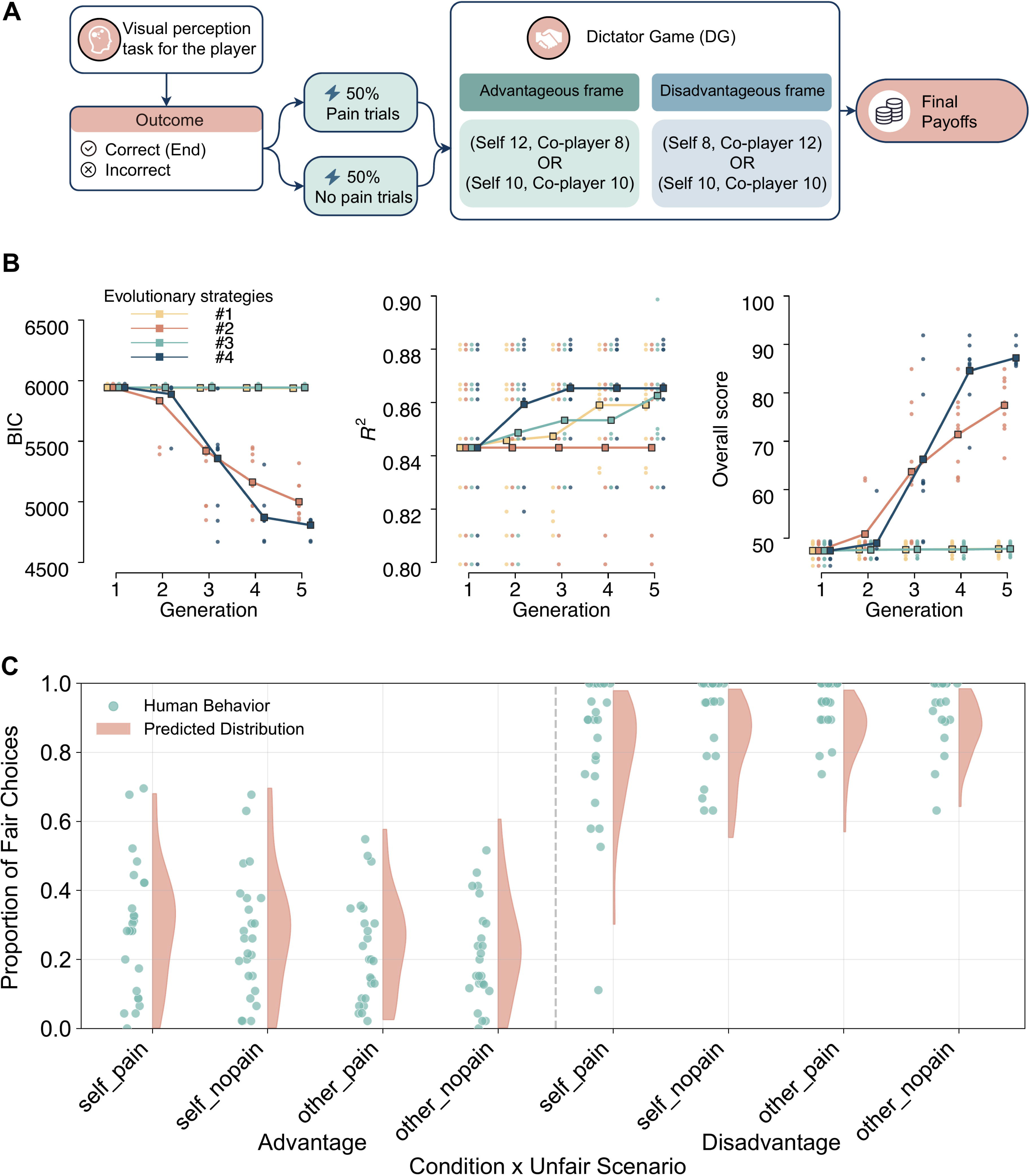
The adapted Dictator Game tested for modeling socioeconomic preferences and evolution dynamics. **A.** Procedure of the Dictator Game under social context (adapted from Gao et al., 2018). In each trial, the participant (the proposer) is paired with an anonymous recipient. Either the participant or the recipient performs a visual perception task. Failure of the task may trigger an electric shock delivered to the recipient, resulting in four social contexts (self/other failure pain/no pain). Subsequently, under the specific social context, the participant chooses between a fair option, in which both players earn 10 tokens equally, and an unfair option, in which the participant earns more (advantage condition) or the partner earns more (disadvantage condition). **B.** Comparison of evolutionary strategies in the ablation study. Dynamics of metrics over generations of the models during the autonomous evolution process using Deepseek R1 (250528) as an example. The curves indicate the average scores (BIC score, parameter identifiability score, and overall score) of the top 10 models in the model base over iterations from the four evolution strategies (Methods; Table 4). Data points indicate the score of each individual model. **C.** Model fitting on the Dictator Game. The predicted distributions by the best human-evaluated computational model (evoluted from Qwen3-235B-A22B) (violin plots) closely matched the participants’ behavior (blue dots) across the eight conditions of the experiment (self/other failure pain/no pain 2 Inequity types), indicated by the chosen proportion of the fair options on the y-axis. Models from all LLMs and their results are available online (see Data Availability Statement).

**Figure 3.**
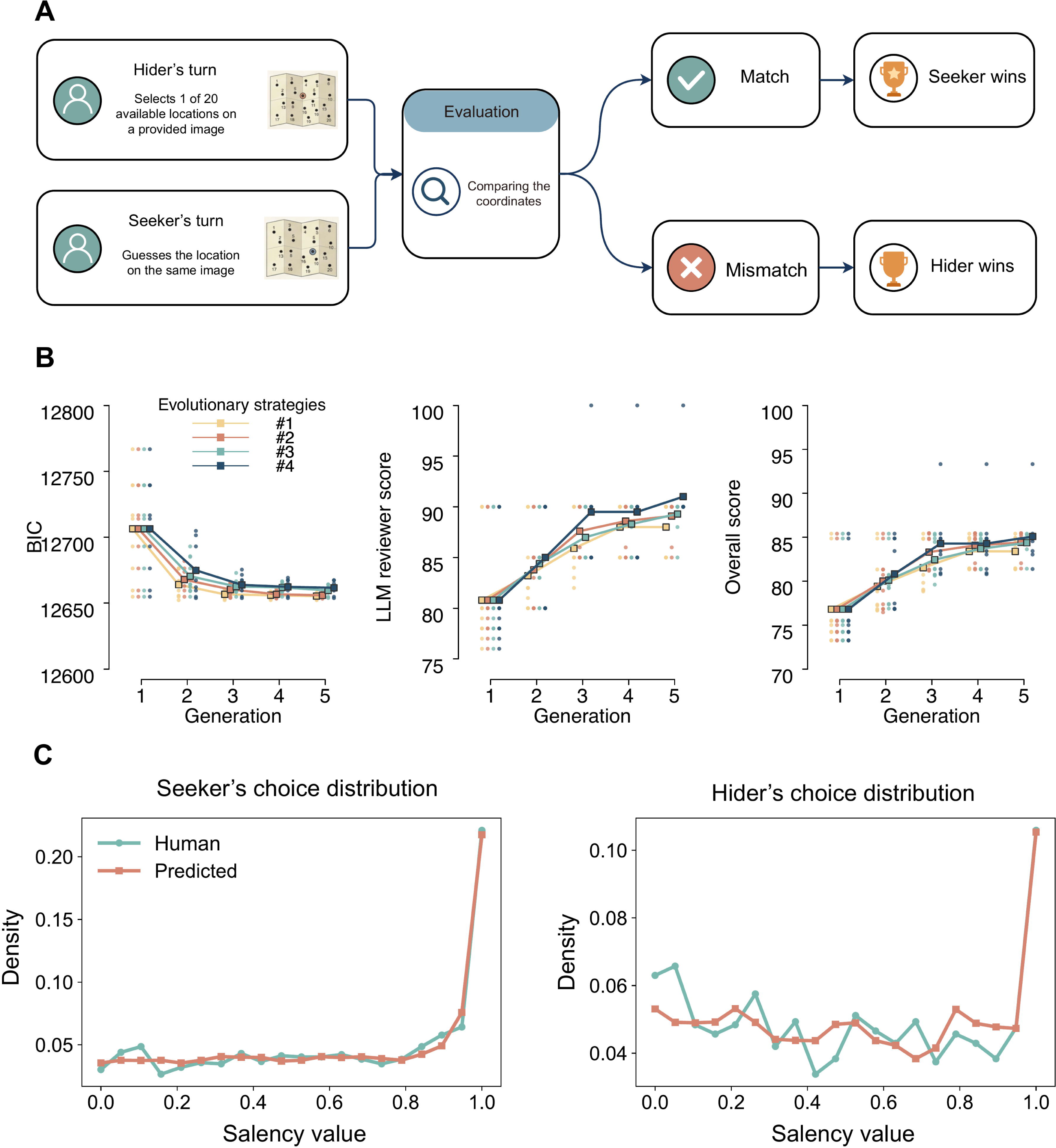
The Hide-and-Seek Location Game for testing theory of mind modeling. **A**. An illustration of the Hide-and-Seek Location Game adapted from Li and Camerer, 2022. The hider and the seeker choose a location on an image of natural scenes. The seeker aims to pick the same location as the hider to win the game; in contrast, the hider aims to avoid being caught by the seeker. **B.** Comparison of evolution strategies in the ablation study. BIC scores reflected the goodness of fit to human behavior, and LLM reviewer scores indicated the models’ interpretability. Overall scores, as a mixture of BIC and LLM reviewer scores, were used for optimization during evolution. **C.** The human-evaluated best model (evoluted from Claude 3.7 Sonnet thinking) predicted the chances of picking a specific location on an image given its saliency score (red lines) closely follow human behavior (blue lines), as the roles of the seeker (left panel) and the hider (right panel).

**Figure 4.**
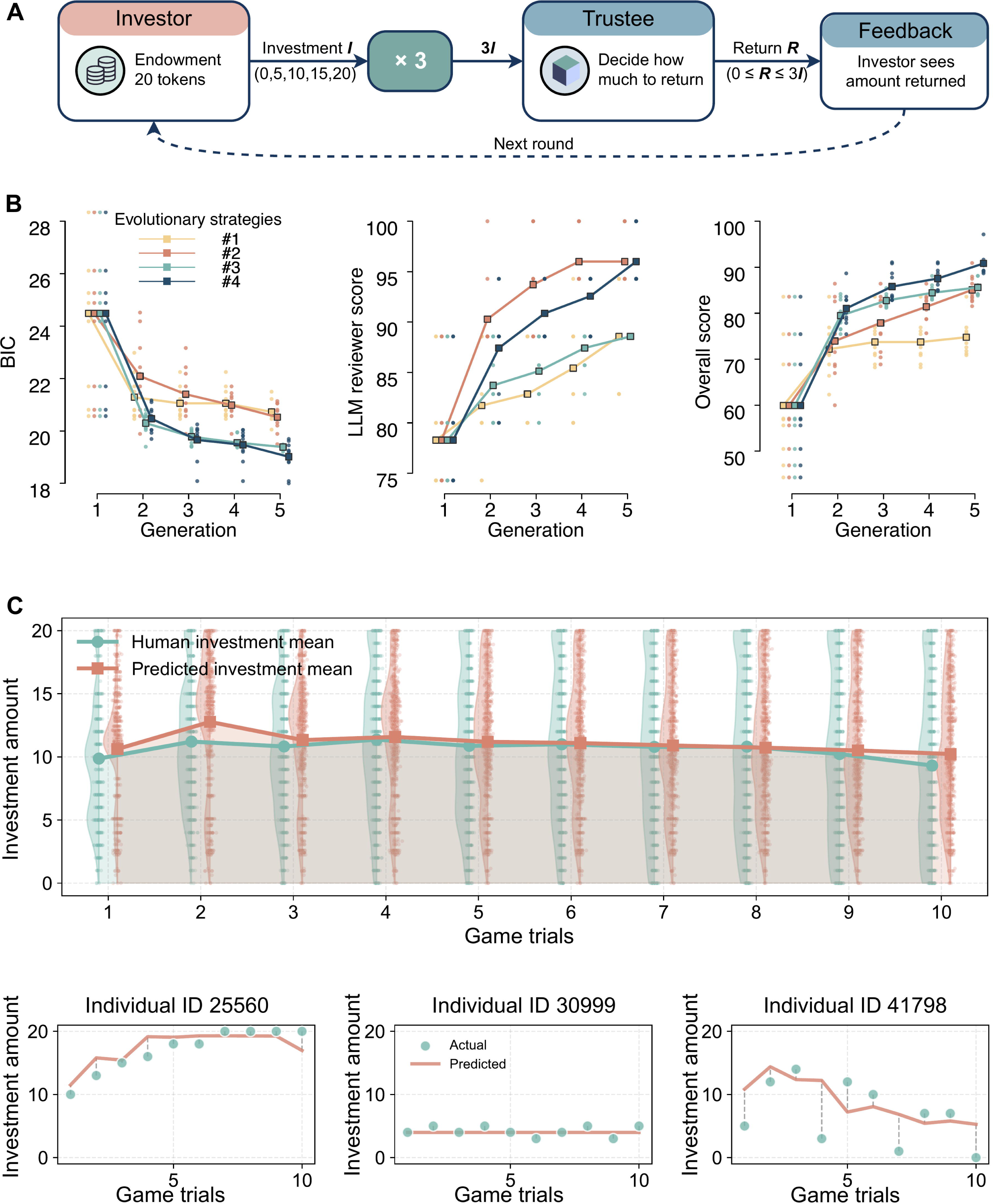
The Trust Game was used to test the interaction between theory-of-mind reasoning and recursive planning over multiple trials. **A.** Illustration of the multi-trial Trust Game. The investor decides to send a proportion of their endowment in each trial of the game to the trustee; the trustee received the investment multiplied by a factor of 3 and decides how much to return to the investor. The same pair of players interact across multiple trials, thereby building reciprocal trust between them. **B.** Performances of four evolution strategies for the Trust Game on metrics of goodness of fit (scored by BIC, left panel), parameter interpretability (provided by the LLM reviewer, middle panel), and the overall score (weighted sum of all metrics and used for optimization during evolution, right panel). **C.** Model fit by the human-evaluated best model (from Gemini 2.5) (green raincloud plots) over 10 consecutive trials of the Trust Game predicted closely the human investment behavior (blue raincloud plots), along with model fits for three representative individuals.

Two human experts in cognitive social science and computational modeling were invited to review the best model from evolutionary strategy #4 from each game. The human evaluators provided detailed feedback and comments written in text, as well as scores on the model’s output over four psychological factors as mentioned.

### The Dictator Game: modeling social preferences and social contexts

The current study systematically evaluated the performance of LLMs across a series of social interaction tasks with varying complexity. The first task we employed was an adapted version of the Dictator Game (Gao et al., 2018), serving as a task with a low complexity level. Unlike the classic Dictator Game, where a proposer is asked to split a fixed amount of resource (e.g., $10) between themselves and a passive recipient, the adapted version consists of binary choices between a fixed fair option (10 vs. 10 tokens) and an alternative unequal option. The unequal options varied to test two distinct types of inequity aversion: advantageous inequity (the proposer earns more than the recipient) and disadvantageous inequity (the proposer earns less than the recipient) (Fig. 2A) (Gao et al., 2018; Sáez et al., 2015).

To further investigate how the LLMs are capable of capturing social contexts and social emotions of humans in impacting their choices, this adapted Dictator Game introduces a preceding perceptual task (dot-estimation) to manipulate the social context dynamically before the binary choices. Specifically, in the perceptual phase, the proposer and the recipient worked as a pair. A mistake in the estimation task could trigger a physical pain stimulation (electric shock) to the recipient. By orthogonally manipulating the identity of the error-maker (Self vs. Other) and the presence of a penalty (Pain vs. No Pain), the 2 × 2 design generates distinct social contexts, most notably an “interpersonal guilt” condition in which one’s own error results in pain to a partner.

In this task, the proposer faces multiple anonymous recipients across independent decision trials. Since there is no feedback or dependency between trials, each decision can be modeled as a standalone event. The objective for the LLM agents was to capture the human proposer’s utility function: specifically, the trade-off between economic preferences (maximizing one’s own reward) and social preferences (sensitivity to the recipient’s outcome and the prevailing social context).

### Autonomous evolution for the Dictator Game

We visualized the evolution results from the ablation study. During the dynamics of autonomous evolution, the goodness-of-fit of the models (indicated by BIC, lower is better, left panel in Fig. 2B) and the parameter identifiability (indicated by *R*^2^, higher is better, middle panel in Fig. 2B) gradually improved and saturated after five generations. Since the LLM reviewer’s interpretability scores were constantly high (at or near the maximum of the scale) for all model candidates in this task due to its low complexity, these scores are not presented. An aggregated overall score, combining goodness-of-fit and parameter identifiability (see Methods), was used as the target function for optimization and visualized in Fig. 2B (right panel).

Comparing across four evolutionary strategies identified Strategy #4, which mixes the parent models with the top-ranked metrics and meanwhile providing the LLM reviewer’s feedback, outperformed the other strategies (see the elaboration on the different strategies in Methods).

The metrics of autonomous evolution from Strategy #4 for all tested LLMs are summarized in Table 1, including a non-reasoning model tested (Deepseek V3) and other reasoning models. Most of the reasoning models performed well with regard to goodness-of-fit. Three of the seven reasoning models achieved lower BIC values than the published human-built benchmark (Gao et al., 2018), and two achieved higher parameter-recovery scores. The non-reasoning model achieved a similar parameter recovery score to the human-built model but a worse goodness of fit than the benchmark. These results show that MindEvolve can generate symbolic candidate models that fit human behavioral data, with several models achieving lower in-sample BIC values or higher parameter-recovery scores than the published benchmark.

**Table 1.**
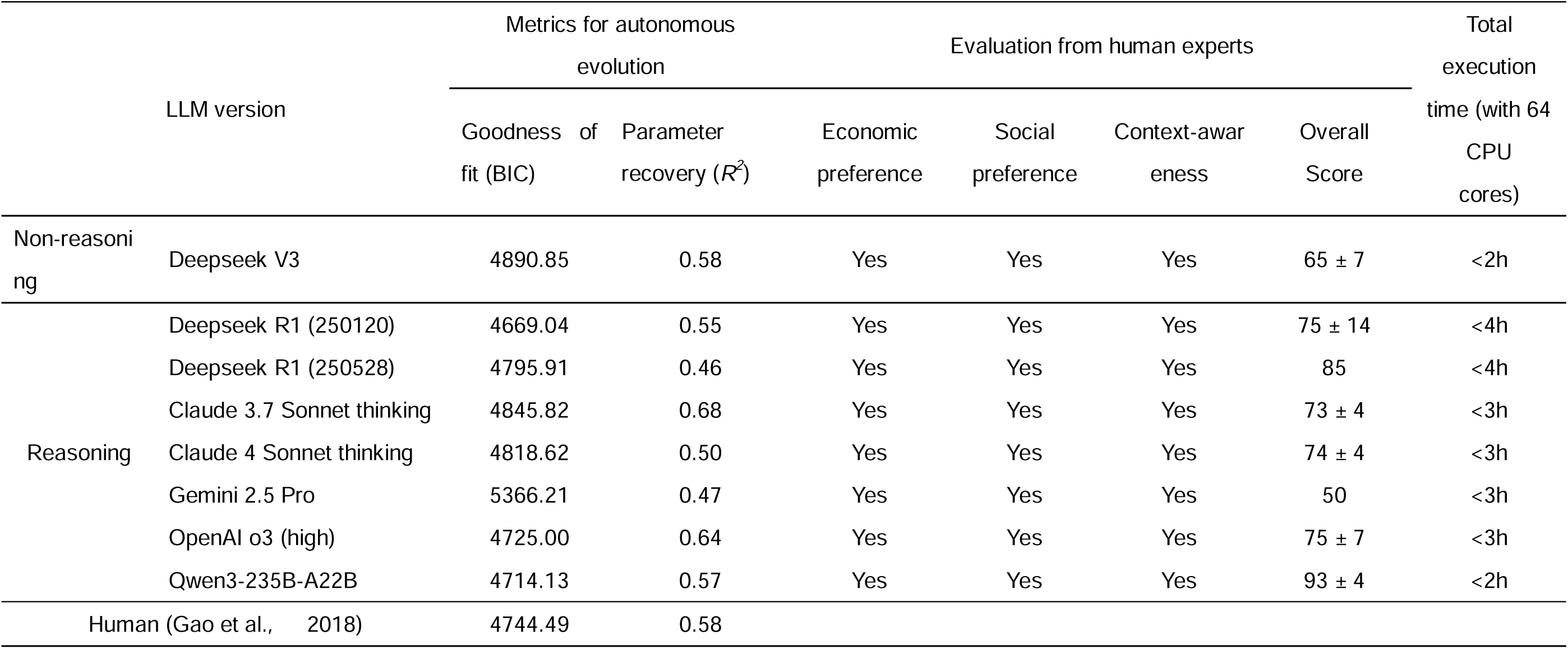
Model performance and human evaluation of LLMs in computational modeling for the Dictator Game.

### Examples of symbolic modeling of the Dictator Game by LLMs

As one of the examples, Claude 3.7 Sonnet thinking reproduced the computational model that was built by the human experts of the reference (Gao et al., 2018). The decision utility of each option can be written as follows (Eq. 1),

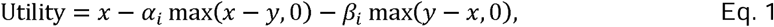

where and indicated the tokens for the proposer and the recipient, respectively, and were used to capture the advantageous and disadvantageous inequality aversion, respectively, with indicating the experimental conditions. Thus, the context modulation is captured by estimating parameters independently across conditions.

The stochastic choice behavior of the human participants is captured by a typical SoftMax function (Eq. 2),

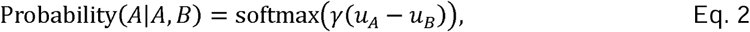

where the utility for the two options (A for the default, fair option, and B for the unequal, alternative option) is calculated according to Eq. 1.

This result indicated that Claude 3.7 Sonnet thinking successfully captures the players’ economic payoff matrices, identifies social preferences by dissociating advantageous and disadvantageous inequality, and accounts for context modulation by estimating parameters separately for each condition.

As another example, the model proposed by Qwen3-235B-A22B captures the key features of the social interaction while additionally parameterizing social-context effects directly on inequity aversion:

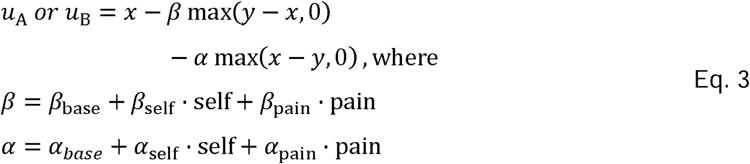

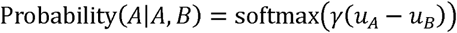

Here *α_base_* and *β_base_* are baseline parameters for advantageous and disadvantageous inequality *α_base_* and *β_base_* capture how responsibility affects fairness preferences, and *α_pain_* and *β_pain_* capture how harm influences fairness preferences. The model evaluates separately advantageous and disadvantageous inequity, weighed by parameters under context modulation. The design of social context parameters can better reflect the interaction between social contextual variables than the human-built model in Gao et al. (2018). The model demonstrates good fit, as shown in Fig. 2C, with the proportion of fair choices by human participants closely matching the model’s predicted distributions across all eight experimental conditions.

### Human-expert evaluation of the Dictator Game demonstrates LLMs’ ability in modeling social preferences with context-awareness

Autonomous evaluation metrics provide quantification and simplified summarization but lose interpretation of the detailed structures of symbolic models. Our developed approach aimed to uncover the detailed structures of the models underlying the generative process, therefore, allowing human evaluation. Two human experts were invited to evaluate the LLMs’ output models evoluted from the best evolutionary strategy - Strategy #4. The Dictator Game involved two basic psychological factors for social interactions, i.e., economic preference and social preference, referring to our framework (Fig. 1). Economic preference was evaluated by whether the model captures the players’ payoff matrices given the description of the game. Social preference was evaluated by players’ psychological concerns beyond their economic payoffs, such as other-regarding and/or inequity-aversion considerations. Crucially, the experimental design included dynamic social contexts to test the LLMs’ capacity for context-awareness as a third factor. It specifically captures how LLMs modulate human’s preferences in response to social emotional changes. Following the findings of Gao et al. (2018), the designed social contexts are critical as they induce distinct psychological states (e.g., guilt) that dynamically modulate human inequity aversion, thus adjust human’s preferences beyond a static baseline. Other than scoring on the three factors, the human experts provided an overall score as a holistic evaluation score of the model. The human-evaluation scores are listed in Table 1.

Overall, all LLMs were successful at capturing the three factors. All LLMs captured the payoff matrices for the players in the Dictator Game, given the game’s simple rule. All LLMs considered the proposer’s social preference, either captured by parameters weighing the recipient’s payoff or aversion parameters to the inequity between the two players’ payoffs. Beyond that, the weighting parameters were adjusted based on the social context. All LLMs well captured the stochasticity of human behavior well using softmax functions.

However, LLMs varied in detailed ways (see SI Appendix, Section S2a for the human experts’ full comments in text), resulting in different overall scores. For example, regarding social preference, most LLMs, such as Deepseek V3, Deepseek R1 (250120), Claude 4 Sonnet thinking, OpenAI o3 (high), did not dissociate between advantage inequity and disadvantage inequity, which, however, has been marked as a key psychological difference in the literature (Gao et al., 2018; Sáez et al., 2015). LLMs such as Gemini 2.5 pro, Claude 3.7 sonnet thinking, Deepseek R1 (250528), and Qwen3-235B-A22B identified the difference between advantage and disadvantage inequity, though Gemini 2.5 pro accounted for the proposer’s aversion to advantageous inequality while neglecting the aversion to disadvantageous inequality.

Regarding the ability of context-awareness, most LLMs considered the social emotional context impact the weighting of the recipient’s payoff. Whereas context modulation was placed on inequity aversions in the model of human-built (Gao et al., 2018). Though these two implementation methods differed only slightly in goodness of fit, the psychological interpretation differ significantly. Interestingly, Deepseek R1 (250528) captured context modulation regarding the recipient’s pain (pain vs no-pain conditions) on the weighting of recipient’s payoff, while placing the responsibility (self-failure and other’s failure) on modulating the advantage inequity. This implementation aligned well with the literature linking empathy to other-regarding concerns (Singer et al., 2006, 2008) and terming the advantage inequity as guilt (Chang et al., 2011; Charness & Dufwenberg, 2006), thus an interesting case that demonstrated the ability of LLMs in integrating a broad range of knowledge.

Human evaluation ranked the model proposed by Qwen3-235B-A22B as the best symbolic model (see the detailed description of the model below), as it dissociated advantage and disadvantage inequity aversion, similar to the human-built model. In addition, it proposed a method for systematically evaluating the impacts of social factors on inequity aversion. Compared with the human-built model, in which the advantage and disadvantage inequity aversion were estimated independently across conditions, this model provided additional insights into the quantification of the impact from the social contexts.

### The Hide-and-Seek Location Game: modeling theory of mind

We tested LLMs’ potential ability to capture the recursive calculation of theory of mind using a game theory task – a spatial version of the classic hide-and-seek game. The Hide-and-Seek Location Game introduces strategic interaction between two players. The hider initiates the game by selecting one location on an image. Images in the game vary by their visual saliency features. The seeker tries to guess the hider’s selection. The hider wins if the seeker fails to reveal the selected location; otherwise, the seeker wins. Though the game is one-shot, the reasoning of the strategies can be recursive and include multiple steps, defined as a level-k problem in game theory (Nagel, 1995; Camerer, Ho, and Chong, 2004, 2015). The hider may choose a hiding spot at random, only influenced by non-reasoning sensory saliency biases originating from the image’s visual features (a level-0 strategy). Instead, they might choose a spot based on their perspective on the seeker’s choice (i.e., a level-1 strategy), instead of guessing randomly (i.e., a level-0 strategy); respectively, the seeker may predict the hider’s perspective on themselves and adjust their strategy accordingly (i.e., a level-2 strategy). The levels of strategic reasoning can increase by many folds until subjects reach their cognitive limit at the k-th iteration. The literature shows that subjects’ behavior can be captured as optimizing against a mixture – or a distribution – of such “level” types, namely, a cognitive hierarchy subjects’ reasoning (Camerer et al., 2004; Li & Camerer, 2022). In the current study, we tested whether LLMs can construct symbolic models that account for higher-order strategic reasoning, specifically the ability to model recursive beliefs and level-k thinking in dyadic interactions, in addition to capturing the specific contexts of the game, such as stimulus-driven attention.

### Autonomous evolution for the Hide-and-Seek Location Game

For the Hide-and-Seek Location Game (Li & Camerer, 2022), we defined two optimization objectives for autonomous evolution: goodness of fit (BIC score) and the LLM reviewer score. A total score combining goodness-of-fit (BIC) and the LLM reviewer score (see Methods) was used for optimization. Fig. 3B visualizes the evolution dynamics of the four different strategies in the ablation study.

The results showed that different evolution strategies exhibited similar dynamics in BIC scores (Fig. 3B, left panel) and Strategy #4 showed an advantage in improving the interpretability of the models as indicated by the faster ramping LLM rreviewer scores (middle panel, Fig. 3B). This result indicated that providing feedback from the LLM reviewer regarding the model interpretability substantially improved the models’ LLM reviewer scores. Strategy #4 achieved the best overall score in improving both goodness of fit and model interpretability after evolution (right panel, Fig. 3B).

### Example model by LLM for the Hide-and-Seek Location Game

The computational model proposed by Claude 3.7 Sonnet thinking provides a comprehensive framework for modeling decision-making in the Hide-and-Seek Location Game by integrating visual perception, the reward structure over the locations, and strategic thinking. The model’s foundation is a cognitive hierarchy model of sophisticated reasoning, where the strategic sophistication levels were assumed to be drawn from a Poisson distribution governed by a role-specific parameter +. At the lowest level (k=0), the choices of both players were assumed to be non-strategic and driven by the visual salience of the locations. The degree of saliency’s impact on behavior was modeled with a power-law transformation of the saliency map to capture every player’s subjective bias.

For higher cognitive levels (k>0), the model introduced a strategic sophisticated reasoning process. A foundation for higher-level reasoning was the explicit modeling of the game’s reward matrices across different locations on the image’s grid, where the seeker aimed to maximize overlap with the hider and the hider aimed to minimize it. A level-k player formed a belief about their opponent’s strategy by taking a weighted average of the opponent’s choices across all levels lower than k. Based on this belief, the player calculated the expected utility of each location. A softmax function, moderated by a rationality parameter, converted these utilities into a choice probability distribution over locations. Eventually, the model posited a dual-process architecture in which each player’s final choice distribution was a weighted mixture of the saliency-driven (level-0) behavior and the aggregated strategic behavior. The visualization of the model’s prediction showed that its behavioral distributions for hiders and seekers as a function of saliency closely match the empirical data (Fig. 3C). Thus, the LLM-built model showed good predictive performance and a clearly defined symbolic model of the theory-of-mind process during social interaction.

### Human-expert evaluation of the Hide-and-Seek Location Game shows success in modeling theory of mind in state-of-the-art LLMs

Two human experts were invited to evaluate the modeling of the Hide-and-Seek Location Game across four dimensions. The Hide-and-Seek Location Game involved one additional component for social interaction than the Dictator Game – theory of mind. That resulted in three dimensions of evaluation from the human experts: a) Task-context awareness was evaluated as whether the model captured the saliency perception of an individual and included it in a manner that biases the individual’s strategies. b) Reward structure was evaluated as to whether the model properly differentiates the utility of their behaviors based on their roles in the game. c) Theory of mind was evaluated as to whether the model captures the iterative process of sophisticated level-k reasoning between the two players. The model code, LLM reviewer comments, and human expert evaluations for these models are available in the Supplementary Materials (see SI Appendix, Section S2b for the human experts’ comments).

Human experts’ evaluations highlighted considerable variation among LLMs (Table 2). Most LLMs captured the task context of visual saliency with respect to human choice strategies. To further capture the payoff matrices based on the two players’ different roles in the game, many models failed to do so, by misrepresenting the payoffs to the different roles. As expected, theory of mind was proved challenging to capture. Many models failed to capture the social reasoning process. A common failure is limiting reasoning to only level-1, responding to the opponent’s strategy but missing the iterative process back and forth between the two players.

**Table 2.**
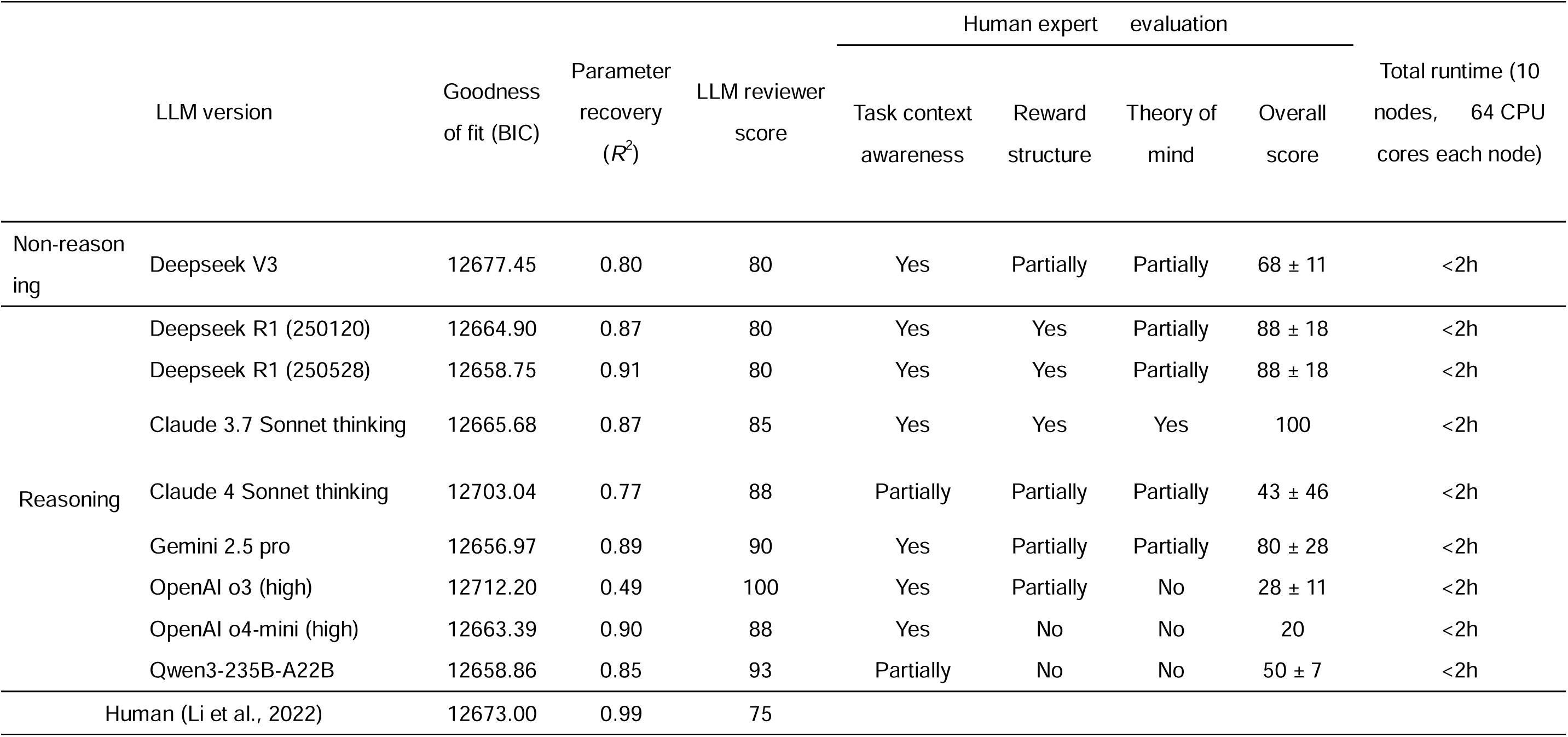
Model performance and human expert evaluation of LLMs in computational modeling for the Hide-and-Seek Location Game.

Nevertheless, Claude 4 Sonnet thinking, Gemini 2.5 pro, and the Deepseek R1 series, were partially successful in implementing the hierarchical reasoning framework with reasonable reward structures. These models allowed agents to form hierarchical beliefs about their opponents’ possible reasoning levels and respond optimally. However, these models still failed to distinguish between the asymmetry between the two roles; instead, they captured the beliefs of both players with the same set of parameters.

Notably, the model generated by Claude 3.7 Sonnet Thinking implemented an explicit cognitive-hierarchy structure relevant to theory-of-mind modeling. The LLM modeled the two players with hierarchical distributions of their beliefs on their opponent’s sophistication levels. Behavioral strategies were inducted based on the visual saliency biases of the two players and optimized iteratively based on the opponents’ sophistication levels. This resulting structure closely resembled the published human-built model (Li & Camerer, 2022).

Results from human evaluation indicated that it remains a significant challenge for most of the tested LLMs to capture the levels of sophistication required for a theory of mind task. Notebly, cutting-edge models of reasoning, such as Claude 3.7 Sonnet thinking, were able to implement a level-k cognitive hierarchy, suggesting that LLM-generated symbolic models can capture more complex human social interactions.

### The Trust Game: modeling theory of mind with recursive planning

Lastly, we moved beyond one-shot games to test a game with multiple trials of social interactions between the same pair of the players, using the multi-trial Trust Game (Hula et al., 2018; Reiter et al., 2023). It further extends the complexity of social interaction by introducing multi-step, reciprocal interaction between two players (Berg et al., 1995; Hula et al., 2018; McCabe et al., 1998). In each trial, the investor (Player 1) receives a fixed endowment and chooses how much (or what proportion) to send to the trustee (Player 2); the sent amount is multiplied by a fixed factor (3× in the tested version), after which the trustee decides how much to return to the investor. The same dyad plays for a fixed number of consecutive trials, and both players observe the full outcome of each trial (investment, multiplied amount, and return), enabling learning and reciprocity to accumulate across trials. Importantly, because future investments can be conditioned on past returns (and vice versa), the game naturally induces reputational incentives and intertemporal trade-offs, making decisions in early trials potentially strategic for shaping later behavior. Compared to the adapted Dictator Game and the Hide-and-Seek Location Game, the Trust Game involves layered social cognition: a) social preferences remain central, but now involve reciprocity (e.g., the trustee returning funds); b) planning requires multi-step reasoning (e.g., the investor forecasting the trustee’s response); and c) theory of mind adds recursive mental-state inference (e.g., “Player 2 believes Player 1 expects generosity”), which is similar to the Hide-and-Seek Location Game. This progression — from one-step to multi-step — introduces significantly increased complexity in social reasoning. It demands theory of mind social reasoning interwined with resursive planning over multiple steps, posing a significant challenge for the LLMs to model.

### Autonomous evolution for the Trust Game

For the Trust Game, we defined two optimization objectives during evolution: goodness-of-fit scores evaluated using BIC and parameter interpretability scores provided by the LLM reviewer. Due to the complexity of the task, parameter recovery could took several months to implement due to its complexity, which was energy-consuming and impractical, thus we left it out from the model evaluation. The overall score as a weighted sum of the BIC scores and the LLM reviewer’s interpretability scores was used for optimization.

Similar to the previous two tasks, the dynamics of evolution from different strategies demonstrated that Strategy #4 achieved the best performance in improving the overall score over the course of evolution (right panel, Fig. 4B). In such a complicated tasks, our designed evolutionary strategies show cleaerer pattern of advantages. Selecting the parent models from the top ranking significantly improved the goodness of fit to the data, as indicated by the lower (better) BIC score curves in Strategies #3 and #4 compared to Strategies #1 and #2 (left panel, Fig. 4B). While providing the LLM reviewer’s interpretability comments as feedback effectively improved the models’ interpretability, as indicated by the faster rising curves on the interpretability scores in Strategies #2 and #4 than in Strategies #1 and #3 (middle panel, Fig. 4B).

### Example model of LLM for the Trust Game

The computational model proposed by Gemini 2.5 pro captures the key elements of the game and integrates psychological factors, such as risk aversion, inequity aversion, theory of mind, and recursive optimization over multiple trials of planning using the Bellman function. First, it establishes correct objective payoffs for both the investor and the trustee. The investor’s payoff is built as the initial endowment subtracted by the invested amount, plus the amount returned by the trustee (Eq. 4),

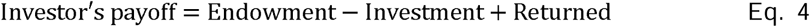

The trustee’s payoff is captured as the multiplied investment subtracted by the amount returned to the investor (Eq. 5),

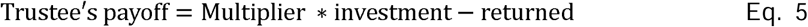

In addition, the investor’s subjective utility is modeled as a combination of economic preference regarding personal gain and social preference regarding the partner’s gain. The economic preference is captured using a power-law function, typically used for capturing risk aversion in the literature (Kahneman & Tversky, 1979); social preference is captured by a weighted term indicating the payoff inequity when the trustee earns more than themselves, i.e., the investor’s disadvantageous inequity aversion which often has a dominant effect in the literature (Eq. 6),

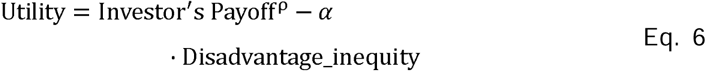

The model integrates these preferences into a strategic planning process using a recursive function that considers both immediate and future utility to maximize outcomes. This planning is integrated with a theory-of-mind component that distinguishes between levels of sophisticated reasoning. At level; 0, the investor forms a Bayesian estimation of the trustee’s return rate based on their past return behavior. At level; 1, the investor engages in perspective-taking, modeling the trustee’s utility by inferring their potential reactions (e.g., guilt or envy) to the investment. However, the implementation does not extend the strategies beyond level 1, it simplifies all levels;k > 1 as equal to level;k = 1.

In summary, the proposed model provides a compelling synthesis of core psychological constructs relevant to the dyadic interaction in the Trust Game. By integrating foundational economic motives with social preferences such as inequity aversion, it yields a rich utility function. The model’s treatment of planning is particularly noteworthy, as it operationalizes strategic, forward-looking decision-making. Furthermore, it offers a concrete, albeit partial, formalization of theory of mind by distinguishing between sophisticated reasoning levels and modeling the crucial act of perspective-taking, thereby capturing key elements of the complex social cognition inherent in the Trust Game.

### Human-expert evaluation of the Trust Game indicates nascent yet promising capability for **complex-task modeling**

Two human experts were invited to evaluate LLMs’ modeling of the Trust Game. The increased complexity of the Trust Game introduced multi-trial planning as an additional dimension to theory of mind and other psychological factors. Thus, it resulted in four dimensions of evaluation in total: a) the economic preference of the players, including capturing of the reward structure of the game and risk taking behaviors when facing uncertainty; b) the social preference of the players, including inequity aversion and other-regarding concerns in addition to their own economic preference; c) theory of mind process capturing the sophisticated reasoning, as in the Hide-and-Seek Location Game; and d) planning recursively over the multiple trials of the game.

The evaluation of different LLMs’ modeling on the Trust Game by two human experts revealed significant disparities in their computational modeling capabilities (Table 3) (see SI Appendix, Section S2c for the human experts’ comments, see the repository for all computational models built by LLMs in Data Availability Statement). Most models correctly specified the players’ economic and social preferences by capturing the game’s payoff structure and incorporating psychological factors such as risk aversion and inequity aversion, which are not captured in the human-built model of the original work (Hula et al., 2018). Again, like in the Dictator Game, this indicated that LLMs perform beyond human-expert in integrate broad range of knowledge in modeling. However, the performance on the factors of theory of mind and planning from most LLMs were scored low. A common flaw was the oversimplification of mechanisms in planning. Instead of recursively optimizing the possible rewards in future steps, many LLMs simplified the future return as additive terms to the current return, such as using temporal discounting over trials.

**Table 3.** Model performance and human expert evaluation of LLMs in computational modeling for the Trust Game.

| LLM version |  | Goodness of fit (BIC) | LLM reviewer score | Human expert evaluation |  |  |  | Overall score | Total runtime (10 nodes, 64 CPU cores each node) |
| --- | --- | --- | --- | --- | --- | --- | --- | --- | --- |
|  |  |  |  | Payoff calculation | Social preferences | Theory of mind | Planning |  |  |
| Non-reasoning | Deepseek V3 | 21.95 | 90 | Yes | Partially | No | No | 35 ± 21 | <3h |
| Reasoning | Deepseek R1 (250120) | 19.23 | 83 | Yes | Partially | No | No | 25 | <4h |
|  | Deepseek R1 (250528) | 22.30 | 87 | Yes | Yes | No | No | 38 ± 18 | <4h |
|  | Claude 3.7 Sonnet thinking | 19.84 | 94 | Yes | Partially | No | No | 31 ± 9 | <6h |
|  | Claude 4 sonnet thinking | 20.53 | 92 | Yes | Partially | No | No | 31 ± 9 | <6h |
|  | Gemini 2.5 pro | 24.68 | 94 | Yes | Partially | Partially | Yes | 90 ± 14 | <160h |
|  | OpenAI o3 (high) | 18.08 | 92 | Yes | Yes | No | No | 40 ± 14 | <10h |
|  | OpenAI o4-mini (high) | 20.93 | 87 | Partially | Partially | No | No | 38 ± 5 | <6h |
|  | Qwen3-235B-A22B | 18.32 | 92 | Partially | Partially | No | No | 30 ± 28 | <3h |
| Human (Reiter et al., 2023) |  | 26.56 | 94 |  |  |  |  |  |  |

**Table 4.**
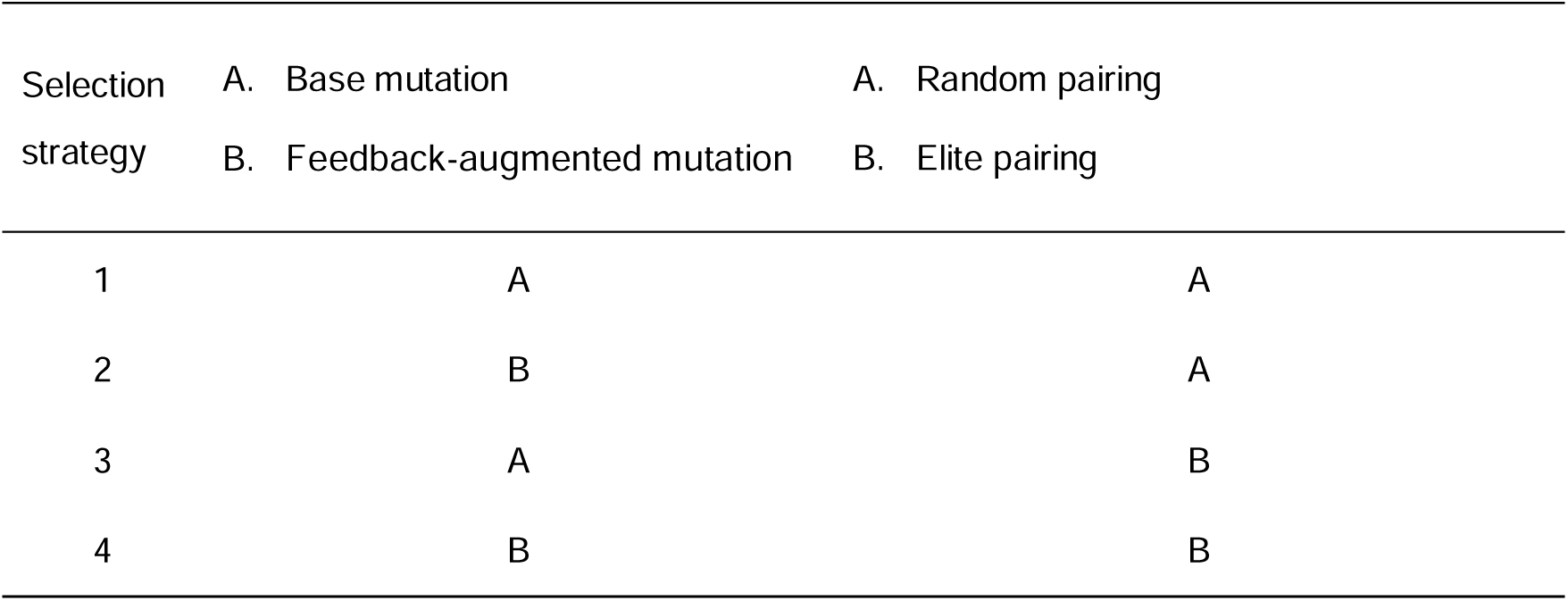
Four evolution strategies were tested for parent-model selection in the ablation study.

| Selection<br>strategy | A. Base mutation | A. Random pairing |
| --- | --- | --- |
|  | B. Feedback-augmented mutation | B. Elite pairing |
| 1 | A | A |
| 2 | B | A |
| 3 | A | B |
| 4 | B | B |

The Trust Game required implementing theory of mind intertwined with planning over trials, since optimizing reward for both the investor and the trustee required estimating the partner’s belief iteratively over trials. This significantly increased the complexity of the modeling task. Most LLMs misinterpreted theory of mind played in the investor as a simple estimation of the trustee’s return rate given their historical return rate, thus, failed in capturing the recursive belief updating central to theory of mind reasoning. Consequently, theory of mind and the dynamic, recursive planning was rarely captured by most LLMs, even though some of these models achieved a better fit to the data (lower BIC scores) than the human-built model. These in-sample differences alone do not establish out-of-sample generalization or superior psychological validity. The result highlights the complementary roles of statistical model-comparison criteria and human-expert evaluation in assessing candidate cognitive models.

Despite these challenges, the model created by Gemini 2.5 pro emerged as a clear top performer, successfully implementing not only economic and social preferences but also recursive planning over trials and a partial formalization of theory of mind (limited to level-1). These findings marked a promising direction of using LLMs to predict human choice behaviors by building interpretable symbolic models, even in tasks with high complexity. The predicted behavior from this model is visualized in Fig. 4C. The three panels at the bottom illustrate three individual examples, which show that the model’s predictions closely fit the observed data.

## Discussion

The current study systematically evaluate the LLMs’ potential in capturing human social interactions via building interpretable symbolic model by human experts on the developed automatic workflow named MindEvolve. We built the workflow using social cognitive science approaches, systematically testing it across four domains of social cognition: the ability to capture the reward structure of agents in task contexts; highly idiosyncratic social preferences; theory of mind and social reasoning; and recursive planning and optimization. With close evaluations from human experts on the symbolic models built by the LLMs, we are able to conclude upon the time of our testing that most LLMs exhibit robust performance in capturing the context-dependent reward structure of the social agents and their social preferences in a one-shot game, e.g., the Dictator Game. When tasked with modeling more complex cognitive processes, such as theory-of-mind-based strategic reasoning and over-trial planning, most LLMs did not achieve reliable performance. However, for a few state-of-the-art LLMs, the generated models implemented theory-of-mind structures that human experts judged to be theoretically relevant. Examples included the model generated by Claude 3.7 Sonnet Thinking for the Hide-and-Seek Location Game and the model generated by Gemini 2.5 Pro for the Trust Game. These findings suggest that future advances in LLMs may enable them to model complex social interactions at a human-expert level. In addition, the LLMs consistently exhibit advantages over human modelers in integrating a broader range of knowledge in their reasoning. The transparency and interpretability of the developed approach through symbolic-model evaluation is key to uncover these findings.

This study has significant theoretical and practical implications. Theoretically, we developed an autonomous workflow to expand LLMs for modeling social interaction, demonstrating their remarkable potential for symbolic modeling in this domain. Our test using social cognitive science approaches and human expert evaluations revealed that certain LLMs can accurately quantify human social and economic preferences and even provide useful insights to discover novel modeling strategies beyond the literature (see modeling for the Dictator Game). For more complex social cognitive mechanisms, most LLMs show limitations in capturing theory of mind and over-trial planning. Using transparent and interpretable strategies, our study effectively revealed these limitations, rather than merely imitating statistical patterns in human behavior (Gurney et al., 2024; Trott et al., 2023). In practice, the built workflow advanced the progress in modeling complex human behaviors. Such an AI-based approach expanded the scope for human researchers to model human social behaviors, which are highly constrained by researchers’ training domains and time-consuming. The developed approach can promisingly enhance the efficiency of traditional research strategies in computational modeling. It may be equally important when considering the advancement of embodied AI. Together, the workflow contributes to “AI for Psychology” by making the generated symbolic models available for direct inspection and expert evaluation, thereby enabling systematic assessment of LLMs’ social interaction modeling in realistic and complex scenarios.

To our knowledge, MindEvolve is the first workflow to apply program-search evolutionary algorithms to systematically test a variety of social interactions. This approach transforms conventional research practices that focus on a limited set of manually designed candidate models by enabling semi-automated generation of numerous high-quality candidate solutions. As a result, symbolic modeling efficiency is greatly improved, and the time required for modeling each task is significantly reduced. The current workflow extends FunSearch (Romera-Paredes et al., 2024) to the domain of social interaction and demonstrates considerable potential for symbolic modeling. Importantly, this study advances beyond CogFunSearch (Castro et al., 2025), whose workflows rely on large-scale iterative optimization involving hundreds of thousands or even millions of LLM calls for program generation and evolution. These advancements enable researchers to tackle increasingly complex tasks by efficiently searching vast spaces that encompass both computational and symbolic models (Castro et al., 2025; Wilson & Collins, 2019), while reducing financial, temporal, and computational resource requirements. Such qualitative improvements shift the trade-off between efficiency and accuracy in computational modeling workflows. MindEvolve thereby establishes an energy-efficient workflow for evolving LLMs in scientific research (Kumar et al., 2024; Mishra et al., 2024).

In computational modeling of the Dictator Game, we found that nearly all LLMs can capture the economic and social preferences of human players and incorporate contextual social factors into their models. Some high-performing LLMs, e.g., Qwen3-235B-A22B and Deepseek R1 (250528), accurately distinguished among the psychological mechanisms underlying advantageous and disadvantageous inequity aversion. Moreover, they can dynamically adjust parameters based on specific contexts and provide fine-grained symbolic representations of variables such as guilt and responsibility attribution. Remarkably, these LLMs can generate novel and reasonable symbolic modeling schemes that have not been proposed in prior work (Gao et al., 2018). These findings indicate that LLMs possess strong knowledge integration and reasoning abilities, enabling them to account for a wide range of factors in modeling, including human risk tolerance and self-interest motives (Cesarini et al., 2009), as well as social preferences for fairness and altruism (Fehr & Schmidt, 1999). Considering social context enables LLMs to model complex social interaction scenarios and demonstrates their potential to be a complementary partner to facilitate human research. However, we also observed that some LLMs fail to distinguish between disadvantageous and advantageous inequity aversion. Differentiating these mechanisms requires a certain level of social cognitive capacity, as the former emerges early in evolution and human development and even in rodents (Watanabe, 2019), whereas the latter is observed only in children with mature cognitive control (typically above age 8; McAuliffe et al., 2017) and also in chimpanzees (Brosnan & de Waal, 2014). Modeling advantageous inequity aversion with LLMs requires understanding advanced social cognition, such as predicting others’ reactions to unfair treatment, recognizing norm violations directed at oneself, and sacrificing immediate gains to maintain long-term cooperative relationships (Brosnan & de Waal, 2014; Gao et al., 2018; McAuliffe et al., 2017).

In the modeling of the Hide-and-Seek Location Game, most LLMs correctly incorporate task context awareness and design reasonable reward structures but fail to model theory of mind. Only Claude 3.7 Sonnet thinking demonstrates a thorough understanding of the underlying logic of level-k and cognitive hierarchy models with limited depth of reasoning. Similarly, in modeling the Trust Game, most LLMs can compute payoffs and subjective utility. However, they generally fail to model theory of mind and planning, revealing that most LLMs are still unable to capture strategies grounded in recursive and iterative social reasoning (Rhoads et al., 2024), particularly when required to infer others’ mental states (Premack & Woodruff, 1978). Nevertheless, we observed some promising results: a few models (e.g., Gemini 2.5 pro) performed relatively well in planning, demonstrating a notable ability to integrate theory of mind into planning, although this captures only a partial aspect of theory of mind in complex social interactions.

These findings indicate that the latest LLMs possess a certain degree of understanding of the planning horizon, as well as the ability to model and interpret planning processes. Planning ability not only affects goal achievement, but also reflects the capacity to consistently execute complex strategies, such as maintaining deception without detection or sustaining stable reciprocal relationships (Hula et al., 2015). Incorporating and estimating planning parameters in LLM modeling allows consideration of the steps humans take to anticipate future outcomes during decision-making. This facilitates precise description and prediction of human behavioral responses in specific contexts and potentially aids in the identification of psychiatric disorders (e.g., ADHD or borderline personality disorder). As Hula et al. (2015) found, individuals with such disorders may display high-level behaviors but are unable to sustain them due to a lack of long-term planning ability.

Regarding theory of mind modeling, results from both the Hide-and-Seek Location Game and Trust Game indicate that current LLMs capture theory of mind at a limited level. They can, to some extent, identify and analyze others’ mental states and beliefs during decision-making and respond to them strategically. However, the level of sophistication is limited. This is consistent with previous findings that LLMs possess a limited understanding of theory of mind (e.g., Kosinski, 2024; Moghaddam & Honey, 2023; Strachan et al., 2024). Their abilities remain at an elementary level and do not meet those of human experts in understanding higher-order theory of mind (Chan et al., 2024; van Duijn et al., 2023; Jones et al., 2024; Kim et al., 2023; Sap et al., 2022). The future development of LLMs may eventually solve this issue and capture the human expert’s level in the aspects of theory of mind reasoning and strategic reasoning. It is important to note that the methods, pathways, and mechanisms employed by LLMs may differ significantly from those of the human brain. This raises concern due to the following reasons. First, LLMs rely primarily on statistical associations from large-scale corpora to “learn” theory of mind, rather than being driven by explicit mechanistic hypotheses (Gurney et al., 2024; Trott et al., 2023). Therefore, LLMs can only generate theory-of-mind-like reasoning within the scope of existing corpora, lacking a deep understanding of underlying cognitive processes and structures, which results in outputs that largely reflect shallow heuristics (Shapira et al., 2024). Second, human abilities to infer others’ mental states depend on rich interpersonal experiences, emotional resonance, and a nuanced understanding of social norms (Frith & Frith, 2006; Koster-Hale & Saxe, 2013). Humans, as active agents (Strachan et al., 2024), can perform multi-level recursive reasoning in complex social environments, a capacity closely linked to coordinated activity in brain regions such as the posterior superior temporal sulcus, temporoparietal junction, and medial prefrontal cortex (Koster-Hale & Saxe, 2013). These regions support the construction of abstract internal models of others’ physical features, mental states, and personality traits, thereby enabling the prediction of others’ behaviors and preferences. In contrast, LLMs may acquire robust pattern recognition abilities through extensive text training; they lack of real-world multimodal information, a physiological basis, and self-awareness. Their performance on theory of mind is more akin to mechanical pattern matching, rather than true “thinking agent” characteristics. Third, unlike human experts, who can systematically decompose complex tasks into actionable submodules and iteratively build and test parameterized models grounded in theoretical foundations, LLMs typically struggle to repeatedly explore diverse possibilities for the same problem. Although the introduction of reasoning mechanisms can partially address this limitation, the depth of LLMs’ reasoning may still fall short of human expert levels.In addition, we found that certain LLMs excelled in specific games: Gemini 2.5 Pro (Trust Game), Claude 3.7 Sonnet Thinking (the Hide-and-Seek Location Game), and Qwen3-235B-A22B (Dictator Game). This phenomenon reflects the “jagged frontier” characteristic of LLM capabilities, whereby LLMs may achieve unexpectedly strong performance on complex or atypical tasks while underperforming on seemingly simpler ones (Dell’ Acqua et al., 2023). One possible explanation is that LLMs, as probabilistic statistical models, are susceptible to biases in training data, inherent cognitive limitations, and limited generalization, which may hinder their ability to comprehensively capture human cognition. These findings highlight the importance of exploring and leveraging the unique strengths of different LLMs. They also suggest that future research should incorporate multiple LLMs into the modeling process, which is particularly crucial for developing general AI agents in dynamic and uncertain real-world scenarios (Gao et al., 2024). Furthermore, appropriate human supervision and adjustment should be considered to address performance challenges arising from “jagged boundaries”.

This study has several limitations to be addressed in future work. First, prior to initiating the automated workflow of MindEvolve, human experts are required to meticulously design the mission prompts, reviewer prompts, and evaluation metrics for specific tasks. In the future, LLMs could assist with prompt generation, and techniques such as fine-tuning may further enhance the capabilities of LLM reviewers, thereby reducing reliance on human experts. Second, the modeling process in this study is knowledge-driven, as LLMs possess extensive prior knowledge. However, LLMs are not well-suited to directly analyzing raw data and can only obtain indirect feedback via the workflow of MindEvolve, which may limit the accuracy of the modeling results. Third, although LLMs perform well in symbolic modeling, they remain susceptible to issues such as hallucination and deception, suggesting that these results should be interpreted with caution.

In conclusion, this study systematically evaluate LLMs’ ability in capturing human behavior during social interactions with interpretable symbolic models by introducing MindEvolve, an autonomous workflow. The high interpretability of this approach facilitates rigorous evaluation from human experts and reveals that while LLMs perform excellently in modeling simple economic and social preferences, most versions of LLMs exhibit limited proficiency in complex social interactions involving theory of mind and recursive planning. A nascent yet promising capability to model complex social interactions has been demonstrated by state-of-the-art LLMs. Our work provide insight for more efficient, autonomous, and interpretable predictions on human social behaviors using AI tools and highlights the current limitations of LLMs in utilizing interpretable mechanisms for complex social cognition.

### Methods Procedure

We implemented MindEvolve (Fig. 1) to conduct computational modeling of social interaction using behavioral datasets from three social-economic games: the Dictator Game (*N* = 37; Gao et al., 2018), the Hide-and-Seek Location Game (*N* = 29; Li & Camerer, 2022), and the Trust Game (*N* = 824; Reiter et al., 2023). The workflow integrated program-search evolution with cognitive-science empowered evaluation and was summarized below.

Prior work on automated model discovery (e.g., FunSearch-based approaches; Castro et al., 2025) provides a foundation for program search. AlphaEvolve (Novikov, 2024) extends this line by introducing an LLM reviewer, explicit feedback for model refinement, and selection mechanisms that integrate multiple evaluation signals during evolution. Building on the AlphaEvolve framework, we implemented an evolutionary algorithm that jointly optimizes goodness-of-fit (BIC), parameter identifiability (parameter recovery *R*^2^), and model interpretability, as assessed by an LLM reviewer. This multi-objective design was aimed at supporting social tasks with high complexity and rich contexts, enabling transparent modeling, and improving search efficiency through parallelized evaluation.

The process began with a task prompt that specifies the modeling objective. The LLM solver was required to explicitly write a symbolic model to predict the player(s)’ behavior in the game by writing an executable Python code. Following the methodology of the source studies (Gao et al., 2018; Reiter et al., 2023), each candidate model was assessed in parallel by an LLM reviewer for interpretability and by a data-fitting program to evaluate the BIC score. We defined a total score (Eq. 7) as a weighted sum of a normalized BIC score (from 0 to 1, lower is better) and a secondary metric M (from 0 to 1, higher is better), which captured the interpretability scores for the location and Trust Games, but model recovery score (*R*^2^) for the Dictator Game. Different indicators were used for the Dictator Game because all proposed models for the Dictator Game achieved high and non-differentiable parameter interpretability due to its simplicity.

More specifically, the BIC scores were linearly mapped to [0, 1] from their original values (from 4,600 to 7,000 in the Dictator Game, from 12,000 to 14,000 in the Hide-and-Seek Location Game, and from 18 to 30 in the Trust Game).

For different games, weights on normalized BIC and M were customized to balance the observed ranges of LLM reviewer score (or *R*^2^), since these values may not span the full range from 0 to 1. Thus, both components contributed comparably to the total score. The total score and the LLM reviewer score were presented as a percentage (out of 100).

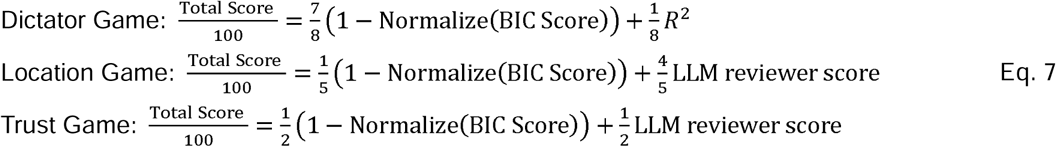

### The autonomous evolution process

LLMs typically generate different computational models over repetitions. Each proposed candidate may perform well along certain dimensions (e.g., goodness-of-fit) and underperform on others (e.g., interpretability). To systematically integrate the strengths of these heterogeneous candidates, we introduced an evolutionary algorithm. In general, the algorithm treated each LLM-generated program as an “individual” in a population and improved the population over generations through selection and mutation. This process is termed “evolution.”

Before evolution, 20 ancestor models were proposed from fixed task-specific prompts and stored in the model base (see the task-specific prompts for the three games in Supplementary Material S1a).

When a new model was proposed, it was evaluated and scored. For model fitting (for all games) and parameter recovery analysis (for the Dictator Game only), we leveraged a high-performance computing cluster comprising 20 servers, each with 128 CPU cores, enabling distributed execution and rapid scoring of large batches of candidate models. For LLM reviewer scores, we designed a dedicated prompt to elicit interpretability ratings from an LLM reviewer (Claude 3.7 Sonnet Thinking was used for all experiments (see the prompt in SI Appendix S1b).

Importantly, we developed evolutionary strategies to pair the parent models to generate new generations. Prior work (Vikhar, 2016) suggests that the choice of evolutionary strategy can substantially affect optimization outcomes. Thus, we compared four evolutionary strategies below.

### Evolutionary strategies

When proposing a new generation, parent models were selected from the model base with different strategies. LLM-based mutation operators were designed to produce offspring models with different strategies (SI Appendix, Section S1c). Together, we designed and compared four evolutionary strategies that differed in parent-model selection and offspring generation (Table 4). Details were listed below.

Strategy #1 (Random pairing + Base mutation): Two parent models were randomly sampled from the model base regardless of their scores. The mutation prompt only contained a plain instruction for the LLM to read both parent programs and generate an offspring program inspired by them.

Strategy #2 (Elite pairing + Base mutation). The parent models were selected from the top 20 ranked by total score. Subsequently, the model with the best BIC was paired with each of the top-10 models ranked by LLM reviewer score, yielding 10 pairs. The model with the highest LLM reviewer score was paired with each of the top-10 models ranked by BIC, yielding an additional 10 pairs. In total, the elite pairing strategy achieved 20 parent pairs. For each pair, we apply the same base mutation prompt as in Strategy #1 to generate an offspring.

Strategy #3 (Random pairing + Feedback-augmented mutation). Parent models were randomly sampled as in Strategy #1. In the generation of the offspring, the mutation prompt was augmented with each parent’s evaluation outcomes, including goodness-of-fit and LLM-reviewer feedback (or parameter recovery score for the Dictator Game), allowing the LLM to condition the generation of offspring on both the parent models and their performance.

Strategy #4 (Elite pairing + Feedback-augmented mutation). This strategy combined the elite pairing as introduced in Strategy #2 and the feedback-augmented mutation prompt introduced in Strategy #3, thereby representing the most delicate design of the evolutionary algorithm.

In summary, Strategies #1 and #3 used random pairing, whereas #2 and #4 adopted elite pairing; Strategies #1 and #2 used the base mutation prompt, whereas #3 and #4 embedded evaluation feedback (goodness-of-fit, LLM reviewer feedback for the Hide-and-Seek Location Game and Trust Game, or parameter identifiability report for Dictator Game) into the prompt. The number of models in the model base accumulated over generations. Since a new generation of the models may improve or degrade relative to their parents, a model proposed in the ealier generation may be selected multiple times as a parent model.

Following a standard machine-learning ablation paradigm (Meyes, 2019), we compared the performance of each strategy for all games (Figs. 2-4). The best models evolved from Strategy #4, the best-performing strategy, were selected for evaluation by the human experts.

## Author Contributions

YY, YF, XZ, FC, BS, and TL conceptualized the research question; YY, YF, BS, and TL designed the test; YY, YF, BS, and TL performed the research; YY, YF, XG, VKD, AH, BS, and TL analyzed portions of the data; and YY, YF, BS, and TL wrote the paper. All authors reviewed and revised the manuscript.

## Data Availability Statement

The supplement materials used in this study are available on the OSF at https://osf.io/p5yrh/overview?view_only=bfc140778c5c4d5ebc95545fc60a99e3. The code and full results are available at https://github.com/MrrDrr/MindEvolve. Data are sourced from previously published studies: Gao et al. (2018), Li and Camerer (2022), and Reiter et al. (2023).

